# Reminders of past choices bias decisions for reward in humans

**DOI:** 10.1101/033910

**Authors:** Aaron M. Bornstein, Mel W. Khaw, Daphna Shohamy, Nathaniel D. Daw

**Affiliations:** Neuroscience Institute and Princeton University, Princeton, NJ, USA; Department of Psychology, Princeton University, Princeton, NJ, USA; Department of Economics, Columbia University, New York, NY, USA; Department of Psychology, Zuckerman Mind, Brain, Behavior Institute, and Kavli Center for Brain Science, Columbia University, New York, NY, USA

## Abstract

We provide evidence that decisions are made by consulting memories for individual past experiences, and that this process can be biased in favor of past choices using incidental reminders. First, in a standard rewarded choice task, we show that a model that estimates value at decision-time using individual samples of past outcomes fits choices and decision-related neural activity better than a canonical incremental learning model. In a second experiment, we bias this sampling process by incidentally reminding participants of individual past decisions. The next decision after a reminder shows a strong influence of the action taken and value received on the reminded trial. These results provide new empirical support for a decision architecture that relies on samples of individual past choice episodes rather than incrementally averaged rewards in evaluating options, and has suggestive implications for the underlying cognitive and neural mechanisms.

---

How do we use experience to guide our choices? One approach is to use a simple heuristic: do what has worked best recently. This idea is captured in prominent reinforcement learning (RL) models, which build a running average of the rewards received for each action taken, updating this average incrementally as new rewards are experienced. Such averages can be updated by error-driven learning, tying this approach to observations of reward prediction errors in the brain [1, 2]. This approach is effective and efficient in the sorts of tasks typically used to study reward learning, which usually involve many repeated choices among a small number of options whose only salient difference is how much or how often they are rewarded [2–5]. A running average falls short, however, in more naturalistic settings, where experience is sparse and relevant evidence might concern only a single, or a small number, of individual experiences [6–12].

Here we provide evidence in favor of a more flexible approach, in which choosers draw on memories for individual instances of relevant previous choices and use them to predict how the current decision might turn out. For example, when deciding where to eat dinner, I might recall a particular evening dining at a particular restaurant, and make my decision based on how much I enjoyed the meal I had that night. Making decisions this way allows us to pick and choose the most relevant of previous experiences, rather than relying on static summary representations. The latter would be useless in considering a brand new restaurant, whereas a richer memory representation of each past experience could be used to develop an informed guess, by generalizing from experiences at similar establishments. Along these lines, Plonsky and colleagues [13] showed that a similarity-based approach to selecting past experiences capture a complex pattern of dependencies of choice behavior on rare, but impactful, past experiences, leading to near-optimal performance in a wide range of dynamic choice tasks. In this way, evaluating options by sampling the most similar previous experiences resembles nonparametric and kernel-based methods long employed in statistics and machine learning, which draw on individual samples of raw experience [14, 15]. The effectiveness of these estimation algorithms reinforces recent interest in the idea that humans and animals draw on memory to flexibly compute decision variables at choice time [11, 16–22], rather than relying on precomputed averages. Indeed, recent work has shown that associations formed during even a single experience can affect later choice behavior [23, 24].

We present results from two experiments in support of this hypothesis. First, we develop and test a computational model of sequential choice that estimates values by selecting and evaluating individual memories of past trials. We show that this sampling model is a better fit to trial-by-trial choice sequences than is a standard incremental learning algorithm, even in the sort of repeated choice task previously studied using RL models. In a second study, we directly test the prediction that value estimation draws on memories of individual trials, by demonstrating a causal effect on choices of specific, selected experiences, brought to mind by refreshing memories of individual past decision trials.

## Results

### Experiment 1

We first formally investigated whether a sampling model could provide a better trial-by-trial fit to human choices than standard incremental learning. To test this, we re-analyzed choice and neuroimaging data from a previously published learning study [4]. Twenty participants (14 presented with neuroimaging data previously, six additional behavior-only participants included here) performed a series of choices between four virtual slot machines with time-varying payoffs (Figure 1a,b). Here, we compared a standard incremental learning rule (of the form tested in the original study) to a sampling model that evaluates each option by retrieving the rewards received at individual past choices of that option, stochastically sampled with probabilities given by their temporal recency (see *Methods*). These sampled rewards are averaged to compute a net value for each option.

**Figure 1:**
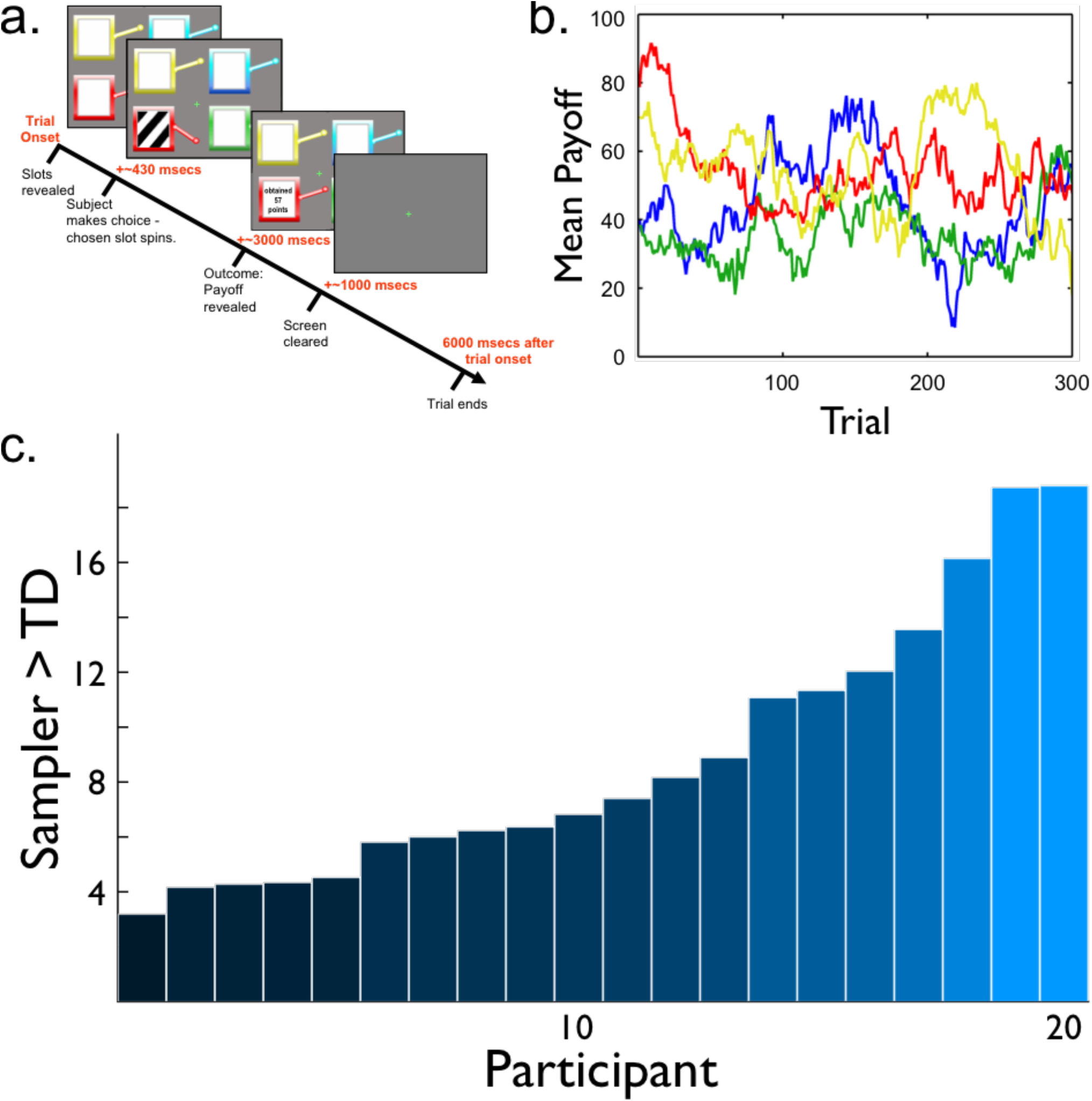
Restless bandit task (from Daw et al., 2006) **a.** Four-armed bandit. Participants chose between four slot machines to receive points. **b.** Payoffs. The mean amount of points paid out by each machine varied slowly over the course of the experiment. **c.** Model comparison. Log Bayes Factors favoring sampling over the TD model.

In the limit, when a potentially infinite number of samples is considered, these sample-based averages would be identical to those computed by the standard model, since the incremental learning rule recursively computes a recency-weighted running average of the experienced outcomes. When fewer samples are considered, the predicted statistics of resulting choices differ systematically. This is because although the value of the sample is, in expectation, the same as the weighted average of the quantities from which it was drawn, individual samples will fluctuate around this mean. For this reason, choices arising from a nonlinear (e.g., max or softmax) comparison of these sampled values across options, do not, in general, have the same statistics as the max of the averages. For instance, the running average approach predicts a fixed, vanishingly small influence of long-past rewards, while in the sampling model events from the far past could still have occasional, but sizable, influence on choices. Furthermore, whereas standard incremental learning models imply that the predicted choice distribution is invariant to the particular sequence of outcomes that gave rise to the options’ values, given only their means, for sampling, even holding constant the average outcomes, the choice distributions will differ depending on the particular sequence (e.g., variance) of outcomes experienced. Our primary model of interest was a sampling model that drew only one sample of past experience. (We also considered variants that drew a greater number of samples, none of these were superior to the one-sample version.) Critically, the two models were carefully set up so as to be matched in all respects other than the key feature of interest (sampling versus averaging): in addition to recency weighting, the choice rules and free parameters were identical.

We computed, for each trial and each participant individually, the probability that each model would produce the choice observed, and estimated the parameter values that maximized these choice likelihoods. Comparing these maximum-likelihood estimates we found that, across the population and for all 20 participants individually, choices were better fit by our sampling model than by the learning model (mean log Bayes Factor against TD 8.8867, SEM 1.0811, exceedance probability > 0.99; Figure 1c). Fit parameters for each model are shown in Table 1.

**Table 1:** Fit model parameters for Experiment 1. The parameters shown are the mean (SEM) across subjects. The final column shows the mean (SEM) of the log Bayes Factor versus the Sampler model (smaller is better).

| Model | $\alpha$ | $\beta$ | $\beta^c$ | log Bayes |
| --- | --- | --- | --- | --- |
| TD | 0.7754 (0.0472) | 8.7069 (0.9564) | 0.6498 (0.0193) | 8.8867 (1.0811) |
| Sampler | 0.7277 (0.0447) | 9.0130 (1.1012) | 0.3046 (0.1288) | - |

We repeated these analyses on simulated data for which the ground truth model was known to verify that these two models were distinguishable from choice behavior (Supplemental notes *Simulated model fits* and *Simulated regression results*, Supplemental Figure S1 and Supplemental Table S1), and also tested the generality of this conclusion under a different specification of the softmax choice noise rule (Supplemental note *Alternative forms of choice noise*). We next examined neural correlates of decision variables from the model (reward prediction error and chosen value), recorded using functional MRI collected on a subset of participants, and verified that, like the choices, both were better accounted for by decision variable timeseries derived from the Sampler model rather than the standard one (Supplemental note *Neuroimaging reanalysis*; Supplemental Figure S2).

### Experiment 2

The above results concord with our hypothesis that human decisions are guided by samples of individual past choices, even in a simple repeated choice task. However, one disadvantage of applying sampling models to this type of task is that trials are essentially similar; we cannot directly observe (and must instead marginalize when computing model fit) which individual trials participants have sampled. For this reason, our conclusions are based on an overall comparison of model fit, rather than a test of clear qualitative features of either model. Moreover, the experiment was correlational, rather than causal. To address this, we performed a second behavioral experiment to provide more direct support for the hypothesis by bringing the sampling process under experimental control, so as to measure the impact of a single, selected experience on choices. We leveraged the fact that multiple aspects of the choice experience can be bound together as a single representation. Therefore we used choice-incidental – but still trial-unique – information to tag each choice as a unique event. Specifically, we modified the bandit task from Experiment 1 so that each trial involved a uniquely identifying photograph of an everyday object – a “ticket” emitted by the chosen slot machine (see also [25]). To simplify the task and analysis, the number of choice options was reduced to two from four, and outcome values were limited to wins or losses of $5 (Figure 2a). The probability that each machine would pay a winning ticket varied from trial to trial (Figure 2b). Other than the ticket presentation, this task matches the sort of “two-armed bandit” traditionally employed in reinforcement learning studies.

**Figure 2:**
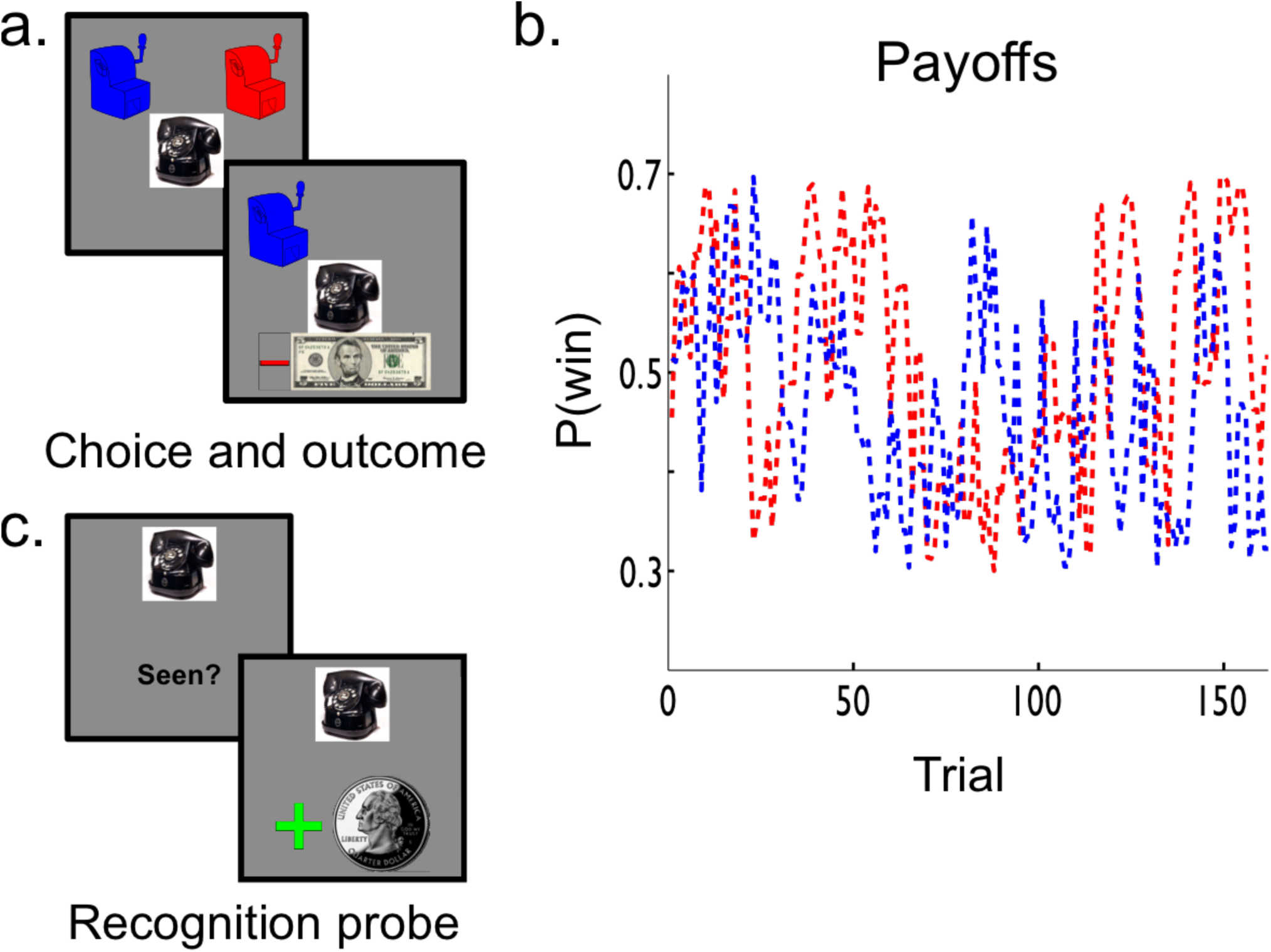
Ticket bandit task. **a.** The ticket-bandit task. Each slot machine (“bandit”) delivered tickets – trial-unique photographs – associated with a dollar value – either −$5 or $5. **b. Payoff probabilities.** The probability of each bandit paying out a winning ticket varied slowly over the course of the experiment. Participants were told that their total payout would be contingent both on the number of winning tickets they accrued and their ability to correctly respond on a post-task memory test asking them to recall the reward value and slot machine associated with each ticket. **c. Memory probes.** Participants encountered 32 recognition memory probes. On 26 of these probe trials, participants were shown objects that were either received on a previous choice trial (“valid”), while on others they were shown new objects that were not part of any previous trial (“invalid”). Participants were asked only to perform a simple old/new recognition judgement – to press “yes” if they had seen the image previously in this task, and “no” if they had not. After each recognition probe, the sequence of slot machine choices continued as before.

Interspersed among the 130 choices were 32 recognition probe trials on which participants were asked whether or not they recognized a given “ticket” image (Figure 2c). These probes were intended to bring to mind the specific trial on which the probed image was first experienced – including the bandit chosen before the ticket appeared and the outcome received after. We hypothesized that this reminder would make the original experience more likely to be sampled during the ensuing choice. In contrast, standard running average models predict no such effect, since they base choices only on the summary statistics. For this reason, this experiment (unlike the previous one) exercises a clear, qualitative prediction between the models: sensitivity to probes evoking individual choices. Probes evoked choices that were, on average, 39 trials in the past – a temporal horizon minimizing both the influence of that reward on a recency weighted running average, as well as the likelihood that the reminded trial would still be present in working memory [26]. After each probe the choice task continued as before. To incorporate reminded trials into the Sampler model, we added a free parameter α_evoked_, specifying the likelihood of reminded trials being incorporated into the sample-based value estimation process. Matching the first analysis, this Sampler model proved a superior fit to choices than did an incremental learning model, for 19 out of 21 participants individually and across the population as a whole, despite being penalized for the additional free parameter (mean log Bayes Factor against TD 6.9182, SEM 1.3227, exceedance probability > 0.99; Figure 3a; model parameters in Table 2).

**Figure 3:**
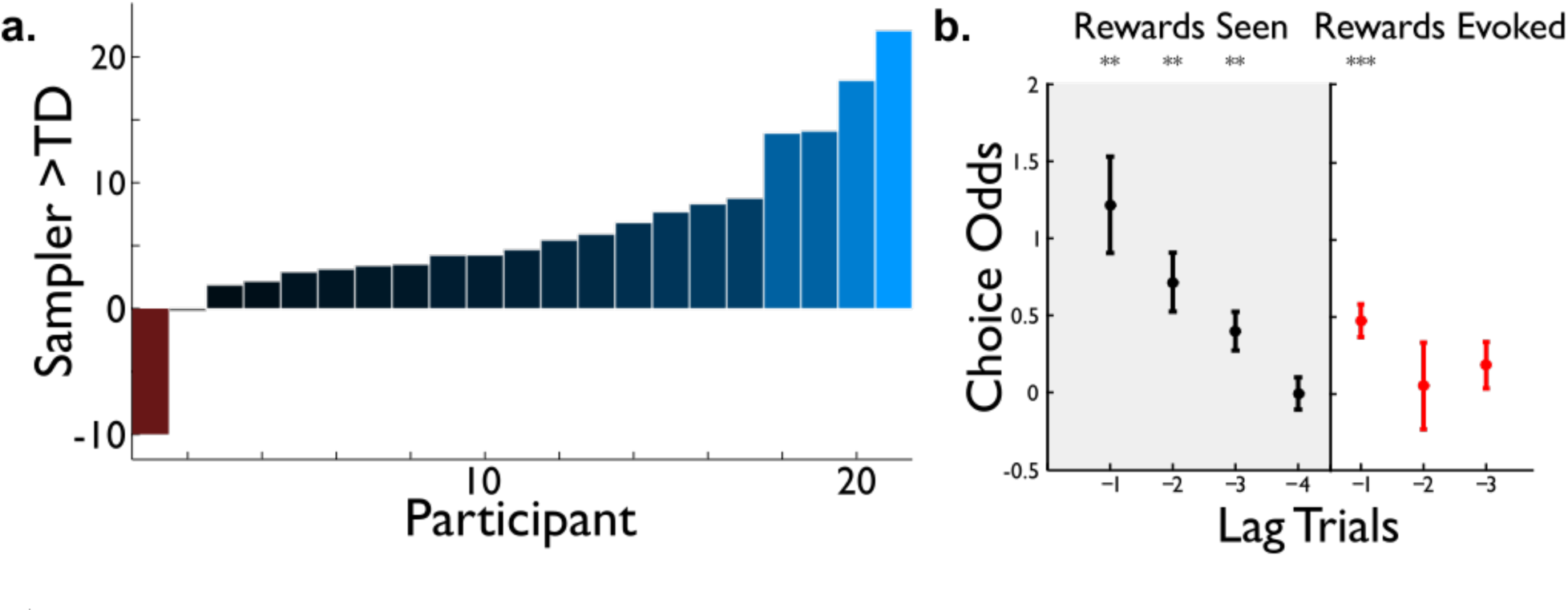
Ticket bandit results. **a. Model comparison.** Log Bayes Factors favoring sampling over the TD model. **b. Impact of probes.** As in standard RL models, choices are affected by previously observed rewards (black points). Here, memory probes evoking past decisions (red) also modulate choices on the subsequent choice trial. Data points are log odds of choosing the righthand option. (* *P* < 0.05, ** *P* < 0.01, *** *P* < 0.001)

**Table 2:** Fit model parameters for Experiment 2. The parameters shown are the mean (SEM) across subjects. The final column shows the mean (SEM) of the log Bayes Factor versus the Sampler model (smaller is better).

| Model | $\alpha$ | $\alpha^{evoked}$ | $\beta$ | $\beta^c$ | log Bayes |
| --- | --- | --- | --- | --- | --- |
| TD | 0.5552 (0.0862) | - | 1.7551 (0.6845) | -0.0930 (0.2354) | 6.9182 (1.3227) |
| Sampler | 0.5393 (0.0583) | 0.4386 (0.0990) | 2.2869 (0.4943) | 0.5855 (0.3215) | - |

However, the key qualitative test of our hypothesis is whether choices following a recognition probe show an effect of the cued trial. For example, if a given recognition memory probe evoked a trial on which the participant chose the blue bandit and was rewarded, then the participant should be more likely to choose the blue bandit on the subsequent choice trial. Conversely, if the choice had resulted in a loss, then they should be more likely to choose the alternative bandit (in this example, red). Incremental RL would be unaffected by the memory probes because it does not maintain memories for individual trials and because the memory probe trials do not provide relevant direct reward experience that could be used to update running average values for the slot machines. Consistent with the sampling model, we observed that choices following a memory probe were also significantly influenced by the much older experience evoked by the probed ticket (*t*(20) = 3.8749, *P* = 0.0009; Figure 3b). The magnitude of the increase in choice probability was comparable to that of a reward directly received between two and three trials in the past, suggesting that the reminder probe was effective in bringing to mind the associated bandit-reward link, and that reward information was then incorporated into choice, on a large fraction of post-reminder trials. Simulations demonstrated that the Sampler model does indeed capture this qualitative pattern of data (Supplemental Figure S1).

## Discussion

These results establish that choices can be made by sampling from individual trial memories, an architecture wholly different from those traditionally envisioned to explain reward learning. We showed that this model can capture the characteristic – and behaviorally observed [27] – recency dependence of incremental averaging approaches, yet provides a superior fit to behavior and decision-related neural signals even in a task originally designed to induce such incremental value updating. This finding goes beyond earlier work that showed that aggregate features of choices – such as a chooser’s variable sensitivity to risky options – are also consistent with a similar sampling model [28, 29]. Critically, the results of Experiment 1 imply that episodic sampling obtains as an influence on choices even when participants are not explicitly encouraged to encode memories of individual trials. However, the results do not directly test the other part of our hypothesis - that choices are driven by memories of individual trials. We therefore tested this idea more directly in a second experiment. In Experiment 2, we strongly encouraged participants to form memories of each trial, so that we could test the influence these memories had on later choices. We brought these past trials to mind by the use of incidental stimulus-stimulus associations present at the time of the original decision, and showed that reward received on that individual trial had a reliable effect on choices following the reminder.

The fact that there was such a strong incentive to encode individual trial memories in Experiment 2 could, in principle, affect the decisions that subjects made. However, just as Experiment 2 extends the results of Experiment 1, Experiment 1 supports the findings of Experiment 2, by showing that the same model obtains on choices in a standard sequential choice experiment (indeed, one that was carried out without reference to the episodic sampling model). Episodic memory is characterized by the formation of stimulus-stimulus associations during a single experience. Therefore this result suggests a key role in sampling decisions for the episodic memory system.

Episodic sampling is a potentially powerful mechanism for choice, as it allows generalizing past experience to apply to novel situations in which decision makers can rely on neither learned action values nor structured, semantic knowledge. In these situations, the episodic memory system could use what features partially overlap with past experience to pattern complete and make predictions for entirely new outcomes using elements of old ones [10, 12]. Importantly, episodic sampling allows these sorts of generalizations to be made at the time of choice, rather than needing to rely on links inferred before the decision at hand [30]. While the latter sort of generalization during encoding can account for many putatively “model-based” generalization phenomena (e.g. [10]), retrieval of episodes or semantic information at the time of a choice is necessary for evaluating unanticipated options [12] and to make use of additional information learned between initial encoding and choice [22]. Such an ability to flexibly generalize – as in latent learning and other revaluation tasks – has previously been associated with a “model-based” learning system, distinct from the “model-free” reward averaging approach [6]. However, although a common assumption is that such behaviors arise from more semantic representations – like cognitive maps – learned over multiple experiences [8], it is possible that samples of individual episodes (e.g. spatial trajectories) might play an analogous role. Thus it remains unclear whether episodic sampling like that considered here supports model-based learning in other tasks, or if it represents yet a third way of deciding [18, 22]. Suggestively, we have previously shown that BOLD signal in the hippocampus at the time of choice scales with the difficulty of making these sorts of “latent learning” decisions – consistent with a sampling process that draws on a wider range of memories when associative information is less decisive about the right action to take [12]. Relatedly, we have shown that enhanced hippocampal-striatal functional connectivity in a bandit task is associated with reduced influence of incremental reward learning on striatal reward prediction errors and choices, and with enhanced episodic encoding for choice options [25].

One open question is the similarity or priority function by which samples are selected from episodic memory. In the terms of nonparametric statistical models, this function corresponds to the kernel, and may itself be modifiable by experience [22]. Here, for simplicity and because of the relative lack of structure in the task, we assumed that samples were drawn according to the recency of the experience. Hertwig and Erev [31] have shown this is a reasonable approximation when the underlying similarity structure of experiences is unknown. But this recency may only be an approximation, and, in environments with more definite structure than the one we use here, different functions or additional dimensions of similarity may obtain. (Indeed, the selection function shown here is likely to be an ineffective learner of tasks in which simple temporal similarity is violated, such as those with payoff rules that alternate from trial to trial.) For instance, Plonsky and colleagues [13] showed that across several kinds of sequential, binary choice tasks, a model that used the recent sequence of outcomes as a template for selecting past experiences had superior performance to one that selected on temporal recency alone. Connecting these sampling procedures to episodic memory opens up the possibility that these effects, and others, can be tied to well-known features of that memory system.

Indeed, the link between reward-guided choice and episodic memory brings into contact two areas of study with well-developed bodies of computational theory and widespread impacts for cognition more broadly [32], and opens the door for leveraging many other features of episodic memory to explain – and perhaps alter – decisions. Though in the present study we focus exclusively on the direct association between action and reward outcome, as the choice trials contain little other useful information, decisions in the real world take place in contexts laden with associative information that could be of relevance to decisions. It seems reasonable to suspect that adaptive decision-makers can leverage this information to support value estimation. Along these lines, in a follow-up study [33], we provide behavioral and neuroimaging evidence that the context in which a sampled experience was first learned predicts the identity of the next experience to be sampled. Further work will be necessary to understand the way in which the episodic nature of memory samples affects the process by which they are sampled.

A related question is to what extent episodic sampling is separate from “model-free” prediction error mechanisms associated with dopamine and its targets in striatum [1, 2]. The finding here that neural RPE signals are captured by expectations formed using associative memories concords with our previous observation that RPE signals during goal-directed decision tasks were better explained by expectations derived from hippocampally-linked associative information [11], and fit with a series of observations that the NAcc RPE signal reflects a mixture of value signals beyond simple model-free TD learning [9, 11, 34]. One possible unifying explanation for these and the previous findings of model-free RPE is that the systems partly overlap, with samples from memory used to train an average-value representation that guides action selection [35]. In this way, rather than computing sampled values separately, direct and sampled experiences might be mixed freely, buffered through a single prediction error signal and net value store [36, 37]. Notably, reward prediction error signals in striatum are influenced by “model-based” and hippocampally linked stimulus-stimulus information [11, 34] and are disrupted when participants successfully form episodic memories for task-irrelevant incidental material [25], suggesting an ongoing, trial-by-trial interaction. (We fit this sort of hybrid model to the data collected in Experiment 2, but the results were equivocal; see Supplemental notes *Adding evoked trials to the TD model* and *Hybrid Sampler and TD model,* and Supplemental Tables S2 and S3)

Altogether, although we show here that episodic sampling can explain choice behavior and neural signals better than incremental learning alone, normative frameworks support the idea that multiple forms of value learning coexist and their influence on each given choice fluctuates according to momentary demands [6, 38, 39]. This finding concords with those cooperative frameworks by showing that the sampling mechanism may contribute to decision-making continually, not just in situations of low experience or uncertain associative structure (c.f. [18]). A superposition of two recency-weighted processes may be one reason why several studies have observed that actions exhibit a double-exponential form of dependence on past outcomes [27, 40] – with components that have been linked to separate activity in striatum and hippocampus [11, 41]. The results also connect to a known role for episodic memory in ongoing elaborative prospection about future situations [42].

More broadly, an explicit link between episodic memory and adaptive instrumental control may serve as one rationalization for why it is that organisms encode rich, associatively-dense memories of individual events in the first place [43]. An open question is to what degree the content and persistence of certain memories, rather than others, can be attributed to the potential usefulness of those memories for later decision-making.

## Acknowledgements

A.M.B. was supported by National Institute of Mental Health predoctoral research fellowship F31MH095501 and a grant from the John Templeton Foundation (Grant ID #57876). D.S. and N.D.D. acknowledge support by National Institute for Drug Abuse R01DA038891 and National Institute for Neurological Disorders and Stroke R01NS078784, and N.D.D. acknowledges support by a Scholar Award from the James S. McDonnell foundation. The authors wish to thank John O’Doherty, Ben Seymour, Peter Dayan, and Raymond J. Dolan, our collaborators on the study reanalyzed here, Yael Niv for helpful suggestions, and Aubrey Clark-Brown for discovering a bug in an early version of the experiment.

## Author contributions

N.D.D. contributed data from Experiment 1; M.W.K and A.M.B. performed the reanalysis of Experiment 1; A.M.B, N.D.D., and D.S. contributed to conceptualizing a role for episodic memory in decision-making; A.M.B. and N.D.D. designed Experiment 2; A.M.B. ran Experiment 2; A.M.B. analyzed the data from Experiment 2; A.M.B. and N.D.D. wrote the paper, with input from M.W.K. and D.S.

## Methods

### Experiment 1: Four-armed bandit from Daw et al (2006)

Participants completed a four-choice bandit task, choosing in each of 300 trials between four different slot machines and receiving a payoff (between 0 and 100 points) for their choice. The data analyzed comprise choice timeseries for the 14 fMRI participants reported previously [4], and for behavioral analyses are augmented with 6 additional behavioral-only pilot participants collected for (but not reported with) that study. Over the experiment, the mean payoffs for the machines diffused according to independent Gaussian random walks.

For a detailed description of the experimental methods, materials, and previous analyses, see the prior report [4]. This section describes the new analyses we performed.

#### Analysis: Choice behavior

Two distinct types of models were compared in their effectiveness at explaining each participant’s timeseries of choices. The first implemented a temporal-difference learning (TD) approach that kept a running average estimate of action values. The second followed a strategy of sampling from previous experiences to estimate these values at the time of decision.

The *standard TD* model maintains a value *Q_t_^TD^*(*a*) for each option *a*, updating it following each choice of *a* according to the difference, *r* − *Q_t_^TD^*(*a*), between obtained and expected rewards. The amount of update is controlled by a step-size (learning rate) parameter α^*TD*^. Formally, on each timestep we updated the value estimate of each action according to Equation 1:

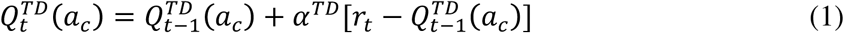

where *r_t_* is the reward received at trial *t*, *a_c_* is the chosen action, and *a_u_* is the unchosen action. Indexing the previous choices of *a* as *i* = 1, 2…*t*-1 choices into the past, this rule can easily be shown [45] at each step to make *Q_t_^TD^*(*a*) a weighted average of the rewards previously received for that choice:

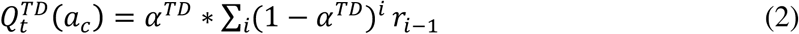

with weights decaying exponentially (by in the choice lag *i*. The resulting decision variable *Q^TD^* is linked to choice probabilities according to the standard softmax function with inverse temperature parameter *β^TD^*. A third parameter, *β^c^*, modeled any perseverative effect of the previous trial’s choice, irrespective of reward received [41, 46]. Choice probabilities were therefore estimated via the combined softmax as:

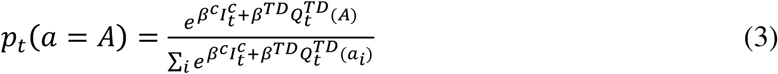

where *I_t_^c^* is an indicator function returning 1 if the previous choice (at trial *t*−1) was identical to that on the current trial (*t*), and 0 otherwise.

A similar sort of temporally decaying dependence on past rewards can also arise from a strategy of recency-based sampling. Instead of maintaining a running average, the sampling model stochastically samples one previous reward for each option with probability given by the same form of weighting:

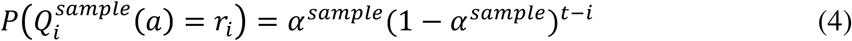

In this model, the most recent experience is most likely to be sampled, and previous trials are successively exponentially less likely to be sampled. (We also considered a version of the model that samples *k* rewards, with replacement, and averages them to produce *Q_t_^sample^*(*a*) this model limits to the standard TD model as *k* → ∞.) The models were equated in all aspects other than the functions they used to aggregate past experience.

For the sampling model, this probability is computed as the likelihood-weighted expectation of Equation 3. One sample is drawn for each bandit at each trial, independently according to Equation 4 within that bandit’s reward history. This expectation is taken over every possible individual sample *r_i_*, with weighting *P*(*Q_i_^sample^*(*a*) = *r_i_*), as given by Equation 4. One sample is drawn for each bandit at each trial, and so the full choice probability is computed as a vector across bandits, over all possible combinations of samples across bandits (four bandits in Experiment 1, and two bandits in Experiment 2). For illustration, in Experiment 2 the computation of *p_i_*(*a* = *A_i_*) is written as Equation 5:

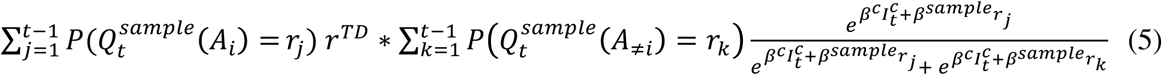

For Experiment 1, the expression is analogous but contains four (rather than two) nested sums, corresponding to the reward sampled for each bandit. In both models, initial conditions (*Q^TD^*(·), *r*_0_) were set to the median payout value (50 for Experiment 1 and 0 for Experiment 2). In the sampling model, any sample probability not assigned to trials *t* = 1, 2… was assigned to *r*_0_.

##### Model comparison

The likelihood of the sequence of obtained choices (each conditional on the rewards and choices up to that point) is then computed as the product over trials of the softmax probabilities, averaging over all combinations of samples in the sampling model, and optimizing the free parameters (*α^x^*, *β^x^*, *β^c^*, with the number of samples *k* = 1 for the sampling models) separately for each participant and model to maximize this likelihood (minimize the negative log likelihood). We compared the fit of candidate models to the choice data using Bayes factors ([47]; the ratio of posterior probabilities of the model given the data). We approximated the log Bayes factor using the difference between scores assigned to each model via the Laplace approximation to the model evidence [48]. In participants for whom the Laplace approximation was not estimable for any model (due to a non-positive definite value of the Hessian of the likelihood function with respect to parameters), we approximated the log Bayes factor using the difference in likelihoods penalized using the Bayesian Information Criterion (BIC; [49]). Model comparisons are reported both on the log Bayes factors as estimated for each individual, and as aggregated across the population. Parameters were estimated using a maximum a posteriori method, accounting for priors over the parameters [48]. The posterior evidence calculations assumed the following prior distributions, chosen to be unbiased over the parameter ranges seen in previous studies [35], and to roll off smoothly at parameter boundaries: for the learning rate parameters, we employed a prior of *Beta*(1.1, 1.1); for the softmax temperatures and perseveration parameters, we employed a prior of *Normal*(0, 10). We also report the “exceedance probability” — the posterior probability that one model is the most prevalent among a set, across a population, as computed using the spm_BMS function included in SPM8 [50].

#### Analysis: fMRI

##### Region of Interest definition

Based on prior studies, we identified regions of interest (ROIs) for each neural decision variable of interest: Chosen Value (CV), in the ventromedial prefrontal cortex (vmPFC), and Reward Prediction Error (RPE), in the nucleus accumbens (NAcc). The NAcc ROI was defined anatomically according to known structural boundaries using the procedure outlined by Breiter and colleagues [51] and used in our previous studies of reward-guided choice; specifically, defining the nucleus’ superior border by a line connecting the most ventral point of the lateral ventricle to the most ventral point of the internal capsule at the level of the putamen [11, 35, 51]. In the case of vmPFC, functional and anatomical boundaries are less well-defined; we defined the current ROI as a 10mm sphere centered on the peak coordinate identified as uniquely responding to “Goal Value” (defined identically to Chosen Value here) by Hare and colleagues [52].

##### Regression analysis

Candidate timeseries for each variable of interest (CV and RPE) were generated according to each of the two models: Temporal-Difference learning (TD), and Sampler. The trial-by-trial predictors were first convolved with a hemodynamic response function for each bandit and then (to account for the hemodynamic lag) we selected the volume two TRs (approximately six and a half seconds) following the relevant event of interest – at decision onset for CV, at outcome receipt for RPE. To account for correlation between the two variables, the timeseries derived from the Sampler model were then orthogonalized against their TD counterparts. Both variables were then entered into a simultaneous linear regression, which computed their relative contribution to explaining the timeseries extracted from each of the ROIs. The design matrix also contained twelve participant motion regressors of no interest. The resulting regression weights were treated as random effects and tested against zero across the population by two-tailed t-test.

### Experiment 2: Ticket bandits

#### Participants

Thirty individuals (one left-handed; 14 female; ages 18–38, mean 23.0) participated in the study. Participants were recruited from the New York University community as well as the surrounding area and gave informed consent in accordance with procedures approved by the New York University Committee on Activities Involving Human participants. All had normal or corrected-to-normal vision. All participants received a fixed fee of $5, unrelated to performance, for their participation in the experiment. Additional compensation of between $0 and $18 depended on their performance on the in-task recognition memory probes ($0.25 each of 32 probes), two pseudorandomly-selected choice trials ($5 each), and a post-task recall test. Average total compensation was $12.

#### Exclusion criteria

Data from nine participants were excluded from analysis due to their being unusable, leaving 21 participants analyzed here. Seven participants were excluded due to response biases that indicated they did not attempt to learn the rewards associated with each bandit — specifically, that their choices were more than 90% to either option in either block of the experiment. Two participants were excluded due to below-threshold performance on the in-task memory probes (below the 90% confidence level – or four or more incorrect).

#### Task design

Participants performed a two-armed restless bandit task — a series of 130 choices between two slot machines — with 32 interspersed memory probe trials, for a total of 162 trials. A compulsory rest period of participant-controlled length was inserted after the 81st trial. The experiment was controlled by a script written in Matlab (Mathworks, Natick, MA, USA), using the Psychophysics Toolbox [53]. Prior to the experiment, participants were given written and verbal instructions as to the types of trials, the buttonpresses required of them, the slot machine payoff probabilities, and the rules for determining final payout. Instructions emphasized that there was no pattern linking the content of the ticket photographs to their dollar value or slot machine. Participants were not told that the memory probe trials should have an effect on their choices, nor was any effect implied. They were, however, told that memorizing the content, source (which slot machine the ticket came from), and dollar value of the tickets would impact their total payout after a post-task memory test.

To aid memory, participants were offered a mnemonic strategy. Specifically, they were told that the photographs could be treated as tickets, which could be placed in one of four imaginary “pockets”: left or right, depending on the slot machine, and front or back, depending on the dollar value. During practice, the keypresses required to advance each trial were instructed in the context of this mnemonic strategy (e.g., “Press ‘a’ to put the ticket in your left hand, now press ‘a’ again to put the ticket in your front pocket.”). Participants completed two practice trials before beginning the main experiment.

#### Choice trials

On each choice trial, participants were presented with two slot machines, each on either side of the upper third of the screen (Figure 2a). The slot machines paid out tickets worth either $5 or −$5. Participants were instructed to press a key corresponding to the slot machine they felt had the best chance of paying out a winning (+$5) ticket rather than a losing (−$5) ticket on that trial. They could choose either the lefthand (key “a”) or righthand (key “b”) slot machine. The slot machines were visually identical except for their color (blue and red), and side of the screen, both of which remained fixed throughout the task. The probability of each machine paying a winning ticket changed independently on each trial according to a diffusing Gaussian random walk with reflecting bounds at 30% and 70% (Figure 2b). At each timestep *t*, *π_t_^i^* — the probability that machine *i* would pay out a winning ticket — diffused according to: *π*_*t*+1_^*i*^ = *π_t_^i^* + *v* for each *i*. The diffusion noise ν was selected from a zero-mean Gaussian with standard deviation *σ_d_* = 0.1. The initial payoff probabilities were set to 60% and 40%, with the identity (side, color) of the superior starting bandit psuedorandomly assigned for each participant.

After selection, the unchosen slot machine was covered, and the chosen machine remained alone on the screen for 0.25 seconds. Then, a trial-unique photograph appeared, and remained on the screen until participants again pressed the key corresponding to their chosen slot machine — in the instructions, this corresponded to the mnemonic memory strategy of selecting the left or righthand side “pockets” for your ticket. When the correct key was pressed, a gray box appeared around the photograph, and remained there for at least 500ms, or additional time up to two seconds depending on when the participants pushed the correct key.

At the end of the timeout, the box disappeared and the dollar value associated with the ticket — either −$5 or $5 — was displayed, as a photograph of a $5 bill, with a green “+” or a red “-” to the left of the bill image. The bill photograph remained on the screen — along with the chosen slot machine, and the trial-unique “ticket” photograph — until the participant pressed a key corresponding to the value: either “a” for −$5 or “b” for $5. In the instructions, this was referred to as “putting the ticket in your front or back pocket”, on the side indicated by the slot machine identity. Once the correct key was pressed, the bill photograph was surrounded by a gray box, and remained on the screen for two seconds. After each trial, a blank screen was displayed for an inter-trial interval of two seconds.

#### In-task recognition memory probes

Beginning after the tenth choice trial, 32 memory probe trials were interspersed at pseudorandom intervals throughout the task (Figure 3c). Each probe trial consisted of a single photograph and the question: “Is this your ticket? (yes/no)”. Twenty six of these photographs were chosen pseudorandomly without replacement from the list of previously seen images; these trials are referred to as valid memory probe trials. The remaining six photographs were novel; these trials are referred to as invalid memory probe trials. Participants were instructed to press ‘a’ if they remembered seeing that image, and ‘b’ if they did not remember seeing that exact image before.

Correct responses — ‘yes’ for previously seen images, and ‘no’ for images that were not displayed on a previous bandit trial — were rewarded with $0.25 added to the participant’s total payout. This additional reward was indicated by a photograph of a US quarter with a green ‘+’ to the left. Incorrect responses resulted in $0.25 being deducted from the participant’s total payout, indicated by a red ‘-’ to the left of an image of a US quarter. Memory probe rewards were displayed for two seconds.

Rewards for memory probes accumulated over the course of the entire task, rather than for randomly selected rounds — so the total payout could be reduced or increased by as much as $8.00. Probe images remained on the screen for up to four seconds — if no answer was entered in that time, the trial was scored as incorrect.

#### Post-task recall memory probes

Before the experiment began, participants were instructed to remember as many complete bandit-outcome-ticket triplets as they could. Their memory for these triplets was tested in 21 post-task memory probes.

Post-task memory probes were drawn only from the images tested during the experiment as valid in-task memory probes. This time, participants were queried as to their recall of each detail associated with the presented probe image: which slot machine it came from, and which dollar value it was associated with. Participants were incentivized to answer correctly by the fact that their total payout in the experiment was predicated on their performance in this memory task. Specifically, payouts for winning ($5) tickets were reduced, to $0, if either post-task recall question was answered incorrectly. Payouts for losing (−$5) tickets were increased, to −$2.50, if both post-task recall questions were answered correctly (Table 3). (The asymmetry in the final values of the winning and losing tickets was intended to encourage participants to attend to both tasks; if losing tickets could be improved to $0 by correct memory task responses, then participants would have less incentive to track the payout values of the bandits – in this case, simply choosing the same option on every trial would have a net positive EV; similarly, if incorrect memory responses only reduced winning tickets to some positive value, then there would be less incentive to remember the ticket images.)

**Table 3:** Ticket values are modified by performance on a post-task recall memory probe. After the main slot machine task, “tickets” paid out by the machines were presented to the participant again. The participant was asked to recall two specific details associated with the ticket: the machine that paid it out (left or right), and the value of the ticket (−$5 or $5). To encourage participants to encode the ticket-machine-value triplet, they were told that the final value of the tickets would depend on both the original value of the ticket and the participant’s performance on two post-task recall questions. If they answered either question incorrectly, $5 tickets were modified to be worth $0. If they answered both questions correctly, −$5 tickets were modified to be worth −$2.5. The payout values of each ticket after the memory tests are described in this table. Values altered by the results of the memory tests are highlighted in bold.

| Probe outcome | \$5 ticket | -\$5 ticket |
| --- | --- | --- |
| Correct on both questions | \$5 | -\$2.5 |
| Incorrect on one question | \$0 | -\$5 |

#### Analysis

Our analysis of choice behavior in this task addressed the hypothesis that memory probe images affect choices in subsequent bandit trials, by evoking the rewards received by choices on past bandit trials.

#### Choice kernel regression

To test this hypothesis, we first examined the effect of past trials — both bandit trials directly experienced, and those evoked by valid memory probes — on choices. Specifically, we entered into a logistic regression one regressor for the rewards received on each of the previous ten trials. If a positive reward was received after choosing the right bandit on trial *t* − τ, this was coded as a 1 in regressor τ, element t. If a negative reward was received after choosing the right bandit, this was coded as a -1. These values were flipped for lefthand bandits: -1 for positive rewards, and 1 for negative rewards.

Next, we included ten regressors coding rewards from past choices that were evoked by valid memory probes over the past ten trials. Specifically, if trial t − τ was a valid memory probe that evoked a trial on which the participant received a positive reward for the right bandit, this was coded as a 1 in the regressor τ, element t, and so forth, following the same coding scheme as directly experienced rewards.

We also entered into the regression matching sets of ten regressors each specifying the identity of the bandit chosen on each of the previous ten trials: −1 for left, 1 for right. And, similarly, for the identity of the bandits evoked by any valid memory probes during the previous ten trials.

Choices of the current bandit were entered as the dependent variable, and coded as -1 (for left) and 1 (for right). The resulting regression weights — indicating the degree to which the current choice was influenced by choices and rewards on a given evoked or directly experienced trial in the past— were tested against zero across the population by two-tailed t-test.

The final regression was thus in the following form:

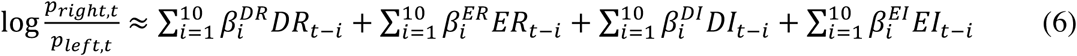

where DR is “directly experienced reward”, ER is “evoked reward”, DI is “directly experienced identity”, and EI is “evoked identity” – each specified over the immediately preceding ten trials.

#### Models

As in Experiment 1, we tested the fit to behavior of computational models capturing our hypothesis and the competing, incremental learning hypothesis.

In the TD model, estimated action values were updated only on bandit choice trials; memory probe trials were not considered to have any impact on the values of the slot machines. This model is identical to Equation 2 from Experiment 1, with the exception that the initial value *Q^TD^*(·) is set to zero, the mean payout on this experiment. In the second model, estimated values were based on one sample drawn from past experience. In this experiment, the sampling model was augmented with an additional parameter, *α^evoked^*, specifying the likelihood that a reminded trial is drawn as a sample. Formally, we distinguish between *direct* and *evoked* rewards *r_i_* according to:

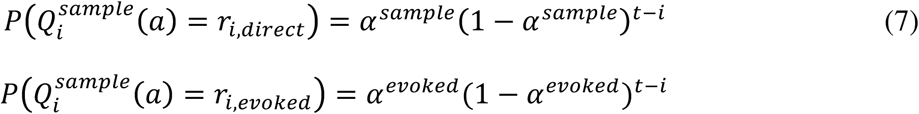

The original Sampling model of Experiment 1, with no impact of reminded trials, is nested within the sampling model, and realized when *α^evoked^* equals zero.)

This formulation was used in computing choice probabilities for all models. These models were otherwise identical to those used in the analysis of Experiment 1.

Models were compared following an identical procedure as used in Experiment 1.

### Data availability

**Data for Experiment 2 are available upon reasonable request to the corresponding author (A.M.B.).**

## References

[1] Andrew C Barto. Adaptive Critics and the Basal Ganglia. In J C Houk, J L Davis, and D G Beiser, editors, Models of information processing in the basal ganglia, pages 215–232. MIT Press, Cambridge, MA, 1995.

[2] W. Schultz, P Read Montague, and Peter Dayan. A Neural Substrate of Prediction and Reward. Science, 275(5306):1593–1599, mar 1997. ISSN 00368075. doi: 10.1126/science.275.5306.1593. URL http://www.sciencemag.org/cgi/doi/10.1126/science.275.5306.1593.

[3] Leo P Sugrue, Greg S Corrado, and William T Newsome. Matching behavior and the representation of value in the parietal cortex. Science, 304(5678):1782–1787, 2004. ISSN 0036-8075. doi: 10.1126/science.1094765.

[4] N D Daw, J P O’Doherty, P Dayan, B Seymour, and R J Dolan. Cortical substrates for exploratory decisions in humans. Nature, 441:876–879, 2006.

[5] Timothy E J Behrens, Mark W Woolrich, Mark E Walton, and Matthew F S Rushworth. Learning the value of information in an uncertain world. Nature Neuroscience, 10(9):1214–21, sep 2007. ISSN 1097-6256. doi: 10.1038/nn1954. URL http://www.ncbi.nlm.nih.gov/pubmed/17676057.

[6] Nathaniel D Daw, Yael Niv, and Peter Dayan. Uncertainty-based competition between prefrontal and dorsolateral striatal systems for behavioral control. Nature Neuroscience, 8(12):1704–1711, 2005.

[7] Alan N Hampton, Peter Bossaerts, and John P O’Doherty. Neural correlates of mentalizing-related computations during strategic interactions in humans. Proceedings of the National Academy of Sciences, 105(18):6741–6746, 2008.

[8] Jan Gläscher, Nathaniel Daw, Peter Dayan, and John P O’Doherty. States versus rewards: dissociable neural prediction error signals underlying model-based and model-free reinforcement learning. Neuron, 66(4):585–95, may 2010. ISSN 1097-4199. doi: 10.1016/j.neuron.2010.04.016.

[9] Dylan A. Simon and Nathaniel D. Daw. Neural Correlates of Forward Planning in a Spatial Decision Task in Humans. Journal of Neuroscience, 31(14):5526–5539, Apr 2011. ISSN 0270-6474. doi 10.1523/JNEUROSCI.4647-10.2011. URL: http://www.jneurosci.org/cgi/doi/10.1523/JNEUROSCI.4647-10.2011.

[10] G. E. Wimmer and D. Shohamy. Preference by Association: How Memory Mechanisms in the Hippocampus Bias Decisions. Science, 338(6104):270–273, oct 2012. ISSN 0036-8075. doi: 10.1126/science.1223252. URL http://www.sciencemag.org/cgi/doi/10.1126/science.1223252.

[11] Aaron M. Bornstein and Nathaniel D. Daw. Cortical and Hippocampal Correlates of Deliberation During Model-Based Decisions for Rewards in Humans. PLoS Computational Biology, 9(12):e1003387, Dec 2013. ISSN 1553-7358. doi: 10.1371/journal.pcbi.1003387. URL http://dx.plos.org/10.1371/journal.pcbi.1003387.

[12] Helen C Barron, Raymond J Dolan, and Timothy E J Behrens. Online evaluation of novel choices by simultaneous representation of multiple memories. Nature Neuroscience, 16(10):1492–1498, sep 2013. ISSN 1546-1726. doi: 10.1038/nn.3515. URL http://www.ncbi.nlm.nih.gov/pubmed/24013592.

[13] Plonsky O, Teodorescu K, and Erev I. Reliance on small samples, the wavy recency effect, and similarity-based learning. Psychological Review, 122(4):621– 647, 2015. doi: 10.1037/a0039413.

[14] Jianqing Fan and Irene Gijbels. Local polynomial modelling and its applications: Monographs on statistics and applied probability. CRC Press, 1996.

[15] Dirk Ormoneit. Kernel-Based Reinforcement Learning. Machine Learning, 49:161–178, 2002.

[16] Ido Erev and Greg Barron. On adaptation, maximization, and reinforcement learning among cognitive strategies. Psychological Review, 112(4):912–31, oct 2005. ISSN 0033-295X. doi: 10.1037/0033-295X.112.4.912. URL http://www.ncbi.nlm.nih.gov/pubmed/16262473.

[17] Neil Stewart, Nick Chater, and Gordon D A Brown. Decision by sampling. Cognitive Psychology, 53(1):1–26, Aug 2006. ISSN 0010-0285. doi: 10.1016/j.cogpsych.2005.10.003. URL http://www.ncbi.nlm.nih.gov/pubmed/16438947.

[18] Mate Lengyel and Peter Dayan. Hippocampal Contributions to Control: The Third Way. Advances in Neural Information Processing Systems, 20:889–896, 2008.

[19] Ian Krajbich, Carrie Armel, and Antonio Rangel. Visual fixations and the computation and comparison of value in simple choice. Nature Neuroscience, 13(10):1292–8, oct 2010. ISSN 1546-1726. doi: 10.1038/nn.2635. URL http://www.ncbi.nlm.nih.gov/pubmed/20835253.

[20] G. Giguere and B. C. Love. Limits in decision making arise from limits in memory retrieval. Proceedings of the National Academy of Sciences, 110(19):7613–7618, apr 2013. ISSN 0027-8424. doi: 10.1073/pnas.1219674110. URL http://www.pnas.org/cgi/doi/10.1073/pnas.1219674110.

[21] Michael Woodford. Stochastic Choice: An Optimizing Neuroeconomic Model. American Economic Review, 104(5):495–500, may 2014. ISSN 0002-8282. doi: 10.1257/aer.104.5.495. URL http://pubs.aeaweb.org/doi/abs/10.1257/aer.104.5.495.

[22] Samuel J Gershman and Nathaniel D Daw. Reinforcement learning and episodic memory in humans and animals: An integrative framework. Annual Review of Psychology, 68:101–128, 2017. doi: 10.1146/annurev-psych-122414-033625.

[23] KD Duncan and D Shohamy. Memory states influence value-based decisions. Journal of Experimental Psychology, General, 145(11):1420–1426, 2016.

[24] Vishnu P Murty, Oriel FeldmanHall, Lindsay E Hunter, Elizabeth A Phelps, and Lila Davachi. Episodic memories predict adaptive value-based decision-making. Journal of Experimental Psychology: General, 145(5):548–558, 2016. ISSN 1939-2222. doi: 10.1037/xge0000158.

[25] G Elliott Wimmer, Erin Kendall Braun, Nathaniel D Daw, and Daphna Shohamy. Episodic Memory Encoding Interferes with Reward Learning and Decreases Striatal Prediction Errors. Journal of Neuroscience, 34(45):14901–14912, 2014. ISSN 1529-2401. doi: 10.1523/JNEUROSCI.0204-14.2014.

[26] Anne G E Collins and Michael J. Frank. How much of reinforcement learning is working memory, not reinforcement learning? A behavioral, computational, and neurogenetic analysis. European Journal of Neuroscience, 35(7):1024–1035, 2012. ISSN 0953816X. doi: 10.1111/j.1460-9568.2011.07980.x.

[27] Brian Lau and Paul W Glimcher. Dynamic Response-by-Response Models of Matching Behavior in Rhesus Monkeys. 84(3):555–579, nov 2005. ISSN http://www.pubmedcentral.gov/articlerender.fcgi?artid=1389781. Journal of the Experimental Analysis of Behavior, 0022-5002. doi: 10.1901/jeab.2005.110-04. URL

[28] Jerker Denrell and James G. March. Adaptation as Information Restriction. Organization Science, 12 (5):523–538, 2001. ISSN 1047-7039. doi: 10.1287/orsc.12.5.523.10092.

[29] Ido Erev, Eyal Ert, and Eldad Yechiam. Loss Aversion, Diminishing Sensitivity, and the Effect of Experience on Repeated Decisions. Journal of Behavioral Decision Making, 21(May):575–597, 2008. doi: 10.1002/bdm.

[30] Sam C. Berens and Chris M. Bird. The Role of the Hippocampus in Generalizing Configural Relationships. Hippocampus, 35(5):591–598, 2017. doi: 10.1002/hipo.22688.

[31] Ralph Hertwig and Ido Erev. The description-experience gap in risky choice. Trends in Cognitive Sciences, 13(12):517–523, 2009. ISSN 13646613. doi: 10.1016/j.tics.2009.09.004.

[32] Daphna Shohamy and Nicholas B Turk-Browne. Mechanisms for widespread hippocampal involvement in cognition. Journal of Experimental Psychology: General, 142(4):1159–70, 2013. ISSN 1939-2222. doi: 10.1037/a0034461. URL http://www.ncbi.nlm.nih.gov/pubmed/24246058.

[33] Aaron M Bornstein and Kenneth A Norman. Putting value in context: A role for context memory in decisions for reward. Nature Neuroscience, in press.

[34] Nathaniel D Daw, Samuel J Gershman, Ben Seymour, Peter Dayan, and Raymond J Dolan. Model-based influences on humans’ choices and striatal prediction errors. Neuron, 69(6):1204–1215, 2011. doi: 10.1016/j.neuron.2011.02.027.Model-based.

[35] Aaron M Bornstein and Nathaniel D Daw. Multiplicity of control in the basal ganglia: computational roles of striatal subregions. Current Opinion in Neurobiology, 21(3):374–80, jun 2011. ISSN 1873-6882. doi: 10.1016/j.conb.2011.02.009.

[36] Richard S. Sutton. Dyna, an integrated architecture for learning, planning, and reacting. ACM SIGART Bulletin, 2(4):160–163, jul 1991. ISSN 01635719. doi: 10.1145/122344.122377. URL http://portal.acm.org/citation.cfm?doid=122344.122377.

[37] Samuel J Gershman, Arthur B Markman, and A Ross Otto. Retrospective revaluation in sequential decision making: a tale of two systems. Journal of experimental psychology. General, 143(1):182–94, 2014. ISSN 1939-2222. doi: 10.1037/a0030844. URL http://www.ncbi.nlm.nih.gov/pubmed/23230992.

[38] Mehdi Keramati, Amir Dezfouli, and Payam Piray. Speed/Accuracy Trade-Off between the Habitual and the Goal-Directed Processes. PLoS Computational Biology, 7(5):e1002055, May 2011. ISSN 1553-7358. doi: 10.1371/journal.pcbi.1002055. URL http://dx.plos.org/10.1371/journal.pcbi.1002055.

[39] Dylan A Simon and Nathaniel D Daw. Environmental statistics and the trade-off between model-based and TD learning in humans. In J Shawe-Taylor, R S Zemel, P Bartlett, F Pereira, and K Weinberger, editors, Advances in Neural Information Processing Systems 24, pages 127–135, 2011.

[40] Greg S Corrado, Leo P Sugrue, H Sebastian Seung, and William T Newsome. Linear-Nonlinear-Poisson Models of Primate Choice Dynamics. Journal of the Experimental Analysis of Behavior, 84(3):581–617, 2005. ISSN 0022-5002. doi: 10.1901/jeab.2005.23-05. URL http://www.pubmedcentral.gov/articlerender.fcgi?artid=1389782.

[41] Aaron M. Bornstein and Nathaniel D. Daw. Dissociating hippocampal and striatal contributions to sequential prediction learning. European Journal of Neuroscience, 35(7): 1011–1023, apr 2012. ISSN 0953816X. doi: 10.1111/j.1460-9568.2011.07920.x. URL http://doi.wiley.com/10.1111/j.1460-9568.2011.07920.x.

[42] Randy L Buckner and Daniel C Carroll. Self-projection and the brain. Trends in Cognitive Sciences, 11(2):49–57, 2006. doi: 10.1016/j.tics.2006.11.004.

[43] J R Anderson. A rational analysis of human memory. In HL Roediger III and F I M Craik, editors, Varieties of Memory and Consciousness: Essays in Honor of Endel Tulving, pages 195–210. Erlbaum, Hillsdale, NJ, 1989.

[44] Hannah M Bayer and Paul W Glimcher. Midbrain dopamine neurons encode a quantitative reward prediction error signal. Neuron, 47(1):129–41, jul 2005. ISSN 0896-6273. doi: 10.1016/j.neuron.2005.05.020.

[45] Brian Lau and Paul W Glimcher. Action and outcome encoding in the primate caudate nucleus. Journal of Neuroscience, 27(52):14502–14, dec 2007. ISSN 1529-2401. doi: 10.1523/JNEUROSCI.3060-07.2007. URL http://www.ncbi.nlm.nih.gov/pubmed/18160658.

[46] R E Kass and A E Raftery. Bayes Factors. Journal of the American Statistical Association, 90(430): 773–795, 1995.

[47] David J C Mackay. Information Theory, Inference, and Learning Algorithms. Cambridge University Press, Cambridge, UK, 2003. ISBN 9780521642989. doi: 10.2277/0521642981.

[48] Gideon Schwarz. Estimating the Dimension of a Model. Annals of Statistics, 6(2):461–464, 1978.

[49] Klaas Enno Stephan, Will D Penny, Jean Daunizeau, Rosalyn J Moran, and J Karl. Bayesian Model Selection for Group Studies. Neuroimage, 46(4):1004– 1017, 2009. doi: 10.1016/j.neuroimage.2009.03.025.Bayesian.

[50] Hans C Breiter, Randy L Gollub, Robert M Weisskoff, David N Kennedy, Nikos Makris, Joshua D Berke, Julie M Goodman, Howard L Kantor, David R Gastfriend, Jonn P Riorden, R. Thomas Mathew, Bruce R Rosen, and Steven E Hyman. Acute Effects of Cocaine on Human Brain Activity and Emotion. Neuron, 19(3):591–611, 1997. ISSN 08966273. doi: 10.1016/S0896-6273(00)80374-8. URL http://www.cell.com/article/S0896627300803748/fulltext.

[51] Todd A Hare, John P O’Doherty, Colin F Camerer, Wolfram Schultz, and Antonio Rangel. Dissociating the role of the orbitofrontal cortex and the striatum in the computation of goal values and prediction errors. The Journal of Neuroscience, 28(22):5623–30, May 2008. ISSN 1529-2401. doi: 10.1523/JNEUROSCI.1309-08.2008. URL http://www.ncbi.nlm.nih.gov/pubmed/18509023.

[52] D H Brainard. The Psychophysics Toolbox. Spatial Vision, 10(4):433–6, Jan 1997. ISSN 0169-1015. URL http://www.ncbi.nlm.nih.gov/pubmed/9176952.

